# Acidic media promotes quiescence entry in *Saccharomyces cerevisiae*

**DOI:** 10.1101/2023.11.20.567958

**Authors:** Alison C. Greenlaw, Toshio Tsukiyama

## Abstract

Quiescence is a conserved cellular state wherein cells cease proliferation and remain poised to re-enter the cell cycle when conditions are appropriate. Budding yeast is a powerful model for studying cellular quiescence. In this work, we demonstrate that the pH of the YPD media strongly affects quiescence entry efficiency in *Saccharomyces cerevisiae*. Adjusting media pH to 5.5 significantly improves quiescence entry efficiency compared to unadjusted YPD media. Thermotolerance of the produced quiescence yeast are similar, suggesting the media pH influences the quantity of quiescent cells more than quality of quiescence reached.

## Description

Quiescence is a conserved process in which cells exit from the mitotic cell cycle for long term survival in eukaryotes. *Saccharomyces cerevisiae* (hereafter yeast) have been leveraged as an effective system to study quiescence, since large quantities of quiescent cells can be readily purified from stationary phase culture (Allen et al., 2006). When quiescent cells are purified using the Percoll gradient method, quiescent (heavy) and non-quiescent (light) fractions are separated, and the percent of cells which enter quiescence can be measured.

We have found that quiescence entry efficiency of the same yeast strain is variable over time, especially dependent upon the batch of YPD (Yeast extract Peptone Dextrose) media used. This was a logistical hurdle, as mutant strains with low quiescence entry efficiency had to be grown and purified at large scale to ensure sufficient quiescent cell yield for genomic and biochemical experiments. Additionally, the necessary scale of the cultures required was difficult to predict due to the variability of quiescence entry. One aspect of the media that was uncontrolled in our laboratory was the pH. Standard recipes for liquid YPD do not include adjusting media pH (“YPD Media,” 2010). Acidic pH has been demonstrated to increase glycerol production (Yalcin & Yesim Ozbas, 2008) and basic pH has been shown to inhibit fermentation and respiration in yeast (Peña et al., 2015). In an industrial strain of yeast, pH 5.5 has been reported as the optimal pH for ethanol production (Narendranath & Power, 2005). We therefore wondered if media pH might influence quiescence entry efficiency.

To test this possibility, three strains in five different pH-adjusted YPD media were grown. Four pH measurements were used: 4.0, 5.56, 6.72 (no adjustment) and 8.24. Since acetic acid is toxic to yeast and promotes aging (Burtner et al., 2009; Murakami et al., 2011), an inorganic acid (hydrochloric acid) and base (sodium hydroxide) were used to test the effect of media pH on quiescence entry. As an additional control, we added low concentration sodium chloride to the 5^th^ YPD, to test whether the pH, or the addition of chloride or sodium ions was responsible for the observed changes to quiescence entry.

Yeast grown in all five batches of media saturated at ∼ 30 OD_660_ per mL, producing approximately 400 total OD_660_ (Figure 1A). Saturated density is essentially equivalent for all five media, with a slight but statistically insignificant decrease for the basic pH media (p = .1410, paired t-test comparing pH = 5.56 and pH = 8.24). In contrast, pH dramatically changes the fraction of cells which are quiescent (Figure 1B). YPD with pH 4.0 results in an approximately 90% quiescence entry efficiency, compared to approximately 25% efficiency at pH 8.24. The addition of 25 mM sodium chloride did not significantly change quiescence entry efficiency compared to unadjusted YPD, demonstrating that pH, not salt ions, is the most relevant variable (p = .7822, paired t-test comparing 25 mM NaCl to unadjusted YPD). Quiescence entry efficiency and pH are highly correlated between pH 4 and pH 8.24 (r^2^ = .9057) (Figure 1C).

**Figure 1.**
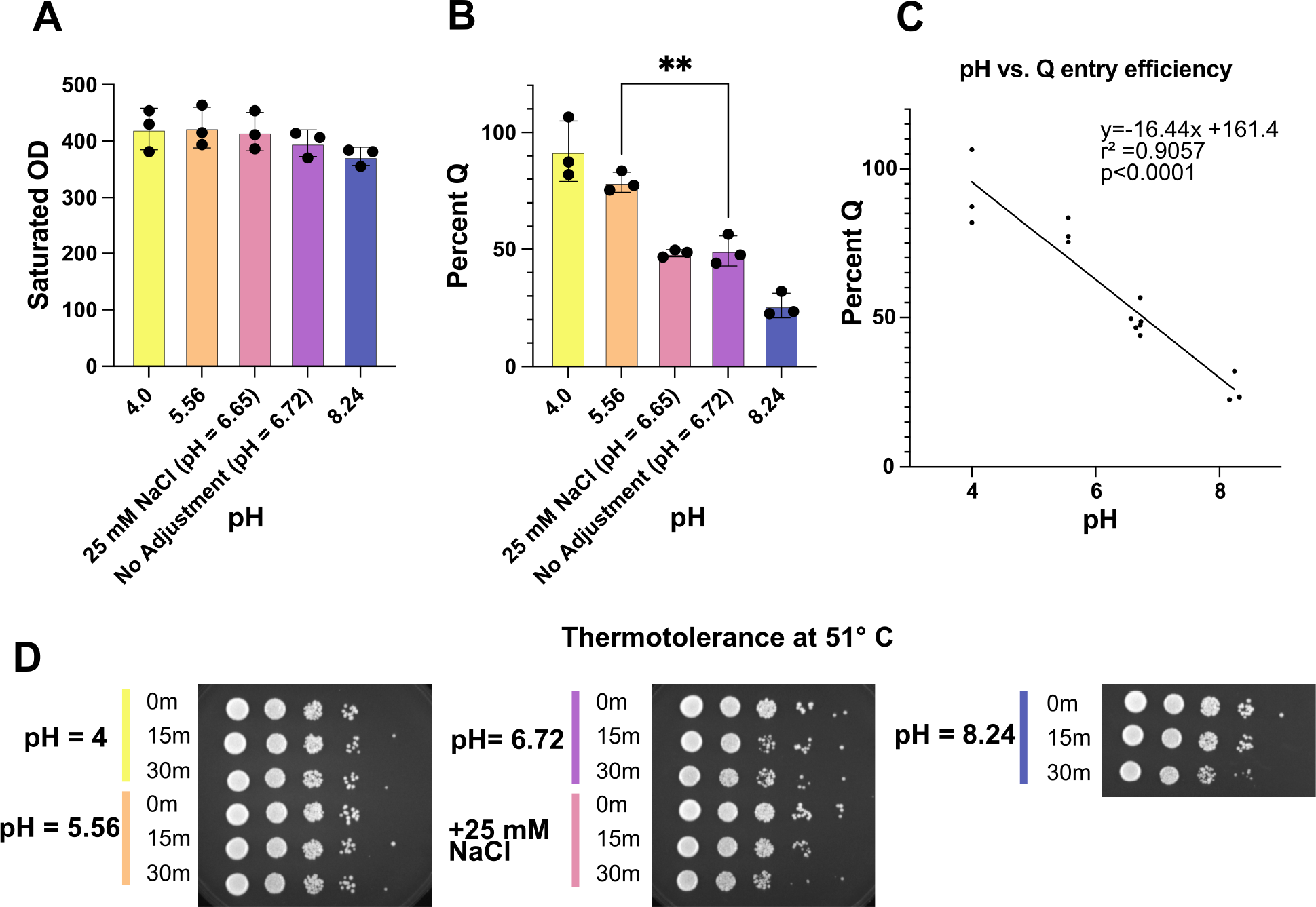
Acidic pH promotes quiescence entry. (**A**) Bar graph showing total saturated OD_660_ of yeast produced by each media condition. Points represent each replicate and bars are standard deviation. Using pair t-test, difference is not significant. (**B**) Bar graph showing percent quiescent cells produced by each media condition. Points represent each replicate and bars are standard deviation. Using pair t-test, difference between 5.56 and 6.72 is significant (** = p < 0.001). (**C**) Scatter plot comparing media pH to percent quiescent. Linear regression analysis performed using Prism. (**D**) Spot test of quiescent cell thermotolerance at 51 °C.

Using simple linear regression, the estimated slope in -16.44, and it is significantly non-zero (p<0.0001). Based on these results, we suggest using YPD media with pH 5.5. We find this results in robust quiescent cells yields and reduces batch to batch variability.

To test if the cells produced were truly quiescent, a thermotolerance assay was performed. Quiescence yeast have improved thermotolerance, and can survive heat shock at 51 °C (Allen et al., 2006; Klosinska et al., 2011). Thermotolerance is relatively similar regardless of media pH (Figure 1D). Although the difference is very minor, pH 4.0 and 5.56 seem to survive slightly better than pH 6.72 and 8.24 after 30 minutes 51 °C. This suggests that the increased quiescence entry yields from the media with pH 4.0 and 5.56 are the result of additional *bona fide* quiescent cells.

In contrast to these results, previous reports demonstrated that acetic acid is toxic to yeast, and low pH has been shown to reduce chronological lifespan (Burtner et al., 2009; Murakami et al., 2011). These experiments were performed in auxotrophic strains, which have poor longevity and are not appropriate for studying quiescence (Boer et al., 2008; Breeden & Tsukiyama, 2022), whereas we used prototrophic strains in our work. Additionally, BY4741 was used for some experiments, which does not enter quiescence (Miles et al., 2023). Therefore, the apparent differences in the effects of low pH media between previous reports and our results are likely due, at least in part, to the differences in strain backgrounds.

## Method

### Making media of different pH

A 2-liter batch YEP was made (1% yeast extract, 2% peptone) and split into five 375 mL aliquots. pH was then adjusted with either HCl (for pH 4 and 5.5) or NaOH (for pH 8.5). As an additional control, NaCl was added at 25 mM to an aliquot. This concentration was determined based on the amount of HCl added to the pH 4 aliquot. The amount of acid, base or salt added was measured, and a remaining volume of water was added to so that 10 mL of additional volume were added to each aliquot. For the unadjusted control, 10 mL of water were added. All 5 media were autoclaved for 1 hour at 120°C. Glucose was added to a final concentration of 2%, and the pH was re-measured. See below table for additional details:

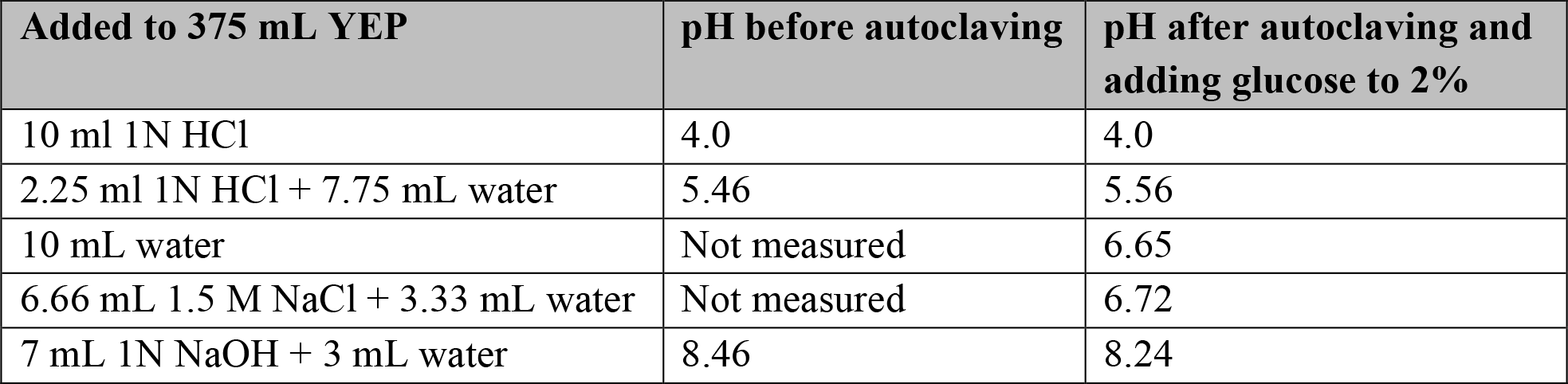

| **Added to 375 mL YEP** | **pH before autoclaving** | **pH after autoclaving and adding glucose to 2%** |
| --- | --- | --- |
| 10 ml 1N HCl | 4.0 | 4.0 |
| 2.25 ml 1N HCl + 7.75 mL water | 5.46 | 5.56 |
| 10 mL water | Not measured | 6.65 |
| 6.66 mL 1.5 M NaCl + 3.33 mL water | Not measured | 6.72 |
| 7 mL 1N NaOH + 3 mL water | 8.46 | 8.24 |

### Quiescent yeast growth, and purification

10 μl of yeast overnight culture from unadjusted YPD was added to each flask and grown for 7 days to saturation at 30 °C on a shaker set to 180 rpm. All cultures were grown in 125 mL flasks with 12.5 mL of media. Saturated density taken by diluting saturated culture 1/100^th^ in YPD and taking OD_660_ using a spectrophotometer. Total saturation density was calculated by multiplying OD_660_ by 100 for dilution and 12.5 for total volume.

Quiescent cells were purified using the Percoll density gradient method as previous described (Allen et al., 2006). Briefly, Percoll and 1.5 M NaCl were combined in an 11:1 ratio, and 12.5 mL of this mixture was added to a gradient column. Columns were centrifuged at 10,000 rpm for 15 minutes at 4 °C to create a gradient. Saturated yeast cells were harvest and resuspended in 3 mL of sterile double distilled water, then the water and yeast mixture was carefully layered over the column. Columns were centrifuged for 1 hour at 1200 rpm at 4 °C. The top of the column was discarded and the bottom ∼ 5 mL section was taken as the quiescent fraction. The quiescent fraction was washed with sterile double distilled water twice and resuspended in 9 mL of sterile double distilled water. Final volume of quiescent fraction was measured using a 10 mL serological pipet.

Quiescent cell yield was taken by diluting quiescent fraction 1/100^th^ in water and taking OD_660_ using a spectrophotometer. OD_660_ was multiplied by 100 for dilution and by the final volume for total quiescent cell yield. Of note, quiescence entry percent was calculated from the total OD_660_ of the saturated culture and total OD_660_ of the quiescent cells. The saturated culture was in YPD and blanked against YPD, while final Q fraction was in water and blanked against water. In one case this led to the total quiescent cell yield being calculated as greater than 100 percent.

### Thermotolerance assay

Heat shock was performed at 51 °C based on previous reports in quiescent yeast (Allen et al., 2006; Klosinska et al., 2011). Quiescent yeast at a concentration 1 OD_660_ per mL were aliquoted into microcentrifuge tubes. Two aliquots were placed in hot water bath at 51 °C for 15 or 30 minutes. An aliquot was kept on the bench as a no heat control. Yeast were serially diluted 1:10 4 times and then 2 μl of each was plated on a YPD plate. The plates were grown at room temperatures for 3 days and then imaged using Biorad Universal Hood II Gel Doc XR System with epi-white light. Images were cropped for the figure to remove empty space, but other image adjustments were performed.

### Statistical Analysis

All experiments were performed in biological triplicate with independently created stains. Graphing and statistical analysis was performed in Prism (version 10). Paired t-test was used to compare between columns in figure 1A and B. Simple linear regression was used to generate the trend line in figure 1C.

## Reagents

| **Strain** | **Genotype** | **Available from** |
| --- | --- | --- |
| yTT5781 | *MATa RAD5+ can1-100* | Tsukiyama Lab |
| yTT5782 | *MATa RAD5+ can1-100* | Tsukiyama Lab |
| yTT5779 | *MAT@ RAD5+ can1-100* | Tsukiyama Lab |

| **Name** | **Supplier** | **Lot number** |
| --- | --- | --- |
| Dextrose (D-Glucose), Anhydrous (Granular Powder/Certified ACS) | Thermo Fisher | D16-10 |
| Gibco™ Bacto™ Peptone | Thermo Fisher | 211820 |
| Gibco™ Bacto™ Yeast Extract | Thermo Fisher | 212750 |
| Hydrochloric Acid | Thermo Fisher | A481-212 |
| Sodium Hydroxide | Sigma Aldrich | S8045 |
| Sodium Chloride | Thermo Fisher | BP538-10 |
| Percoll PLUS | Cytiva | 17544501 |

## Acknowledgements

We thank Dr. Kris Alavattam for advice on statistical tests. We thank Rachel Dell for proofreading.

## References

Allen, C., Büttner, S., Aragon, A. D., Thomas, J. A., Meirelles, O., Jaetao, J. E., Benn, D., Ruby, S. W., Veenhuis, M., Madeo, F., & Werner-Washburne, M. (2006). Isolation of quiescent and nonquiescent cells from yeast stationary-phase cultures. The Journal of Cell Biology, 174(1), 89–100. 10.1083/jcb.200604072

Boer, V. M., Amini, S., & Botstein, D. (2008). Influence of genotype and nutrition on survival and metabolism of starving yeast. Proceedings of the National Academy of Sciences, 105(19), 6930–6935. 10.1073/pnas.0802601105

Breeden, L. L., & Tsukiyama, T. (2022). Quiescence in Saccharomyces cerevisiae. Annual Review of Genetics, 56, 253–278. 10.1146/annurev-genet-080320-023632

Burtner, C. R., Murakami, C. J., Kennedy, B. K., & Kaeberlein, M. (2009). A molecular mechanism of chronological aging in yeast. Cell Cycle (Georgetown, Tex.), 8(8), 1256–1270.

Klosinska, M. M., Crutchfield, C. A., Bradley, P. H., Rabinowitz, J. D., & Broach, J. R. (2011). Yeast cells can access distinct quiescent states. Genes & Development, 25(4), 336–349. 10.1101/gad.2011311

Miles, S., Lee, C., & Breeden, L. (2023). BY4741 cannot enter quiescence from rich medium. microPublication Biology. 10.17912/micropub.biology.000742

Murakami, C. J., Wall, V., Basisty, N., & Kaeberlein, M. (2011). Composition and Acidification of the Culture Medium Influences Chronological Aging Similarly in Vineyard and Laboratory Yeast. PLOS ONE, 6(9), e24530. 10.1371/journal.pone.0024530

Narendranath, N. V., & Power, R. (2005). Relationship between pH and Medium Dissolved Solids in Terms of Growth and Metabolism of Lactobacilli and Saccharomyces cerevisiae during Ethanol Production. Applied and Environmental Microbiology, 71(5), 2239–2243. 10.1128/AEM.71.5.2239-2243.2005

Peña, A., Sánchez, N. S., Álvarez, H., Calahorra, M., & Ramírez, J. (2015). Effects of high medium pH on growth, metabolism and transport in Saccharomyces cerevisiae. FEMS Yeast Research, 15(2), fou005. 10.1093/femsyr/fou005

Yalcin, S. K., & Yesim Ozbas, Z. (2008). Effects of pH and temperature on growth and glycerol production kinetics of two indigenous wine strains of Saccharomyces cerevisiae from Turkey. Brazilian Journal of Microbiology: [Publication of the Brazilian Society for Microbiology], 39(2), 325–332. 10.1590/S1517-838220080002000024

YPD media. (2010). Cold Spring Harbor Protocols, 2010(9), pdb.rec12315. 10.1101/pdb.rec12315

